# Parkinson mutations in *DNAJC6* cause lipid defects and neurodegeneration that are rescued by Synj1

**DOI:** 10.1101/2022.04.27.489745

**Authors:** Julie Jacquemyn, Sabine Kuenen, Jef Swerts, Benjamin Pavie, Vinoy Vijayan, Ayse Kilic, Dries Chabot, Yu-Chun Wang, Nils Schoovaerts, Patrik Verstreken

## Abstract

Recent evidence links dysfunctional lipid metabolism to the pathogenesis of Parkinson’s disease, but the mechanisms are not resolved. Here, we created a new *Drosophila* knock-in model of *DNAJC6/Auxilin* and find that the pathogenic mutation causes synaptic dysfunction, neurological defects and neurodegeneration, as well as specific lipid metabolism alterations. In these mutants membrane lipids containing long-chain polyunsaturated fatty acids, including phosphatidylinositol lipid species that are key for synaptic vesicle recycling and organelle function are reduced. Overexpression of another protein mutated in Parkinson’s disease, Synaptojanin-1, known to bind and synthesize specific phosphoinositides, strongly rescues the *DNAJC6/Auxilin* neuronal defects and neurodegeneration. Our work reveals a functional relation between two proteins mutated in Parkinson’s disease and implicates deregulated phosphoinositide metabolism in the maintenance of neuronal integrity and neuronal survival in Parkinsonism.

## INTRODUCTION

Clathrin-mediated endocytosis (CME) is a vital eukaryotic process controlling the shuttling of cargo and various types of lipids from the Golgi network or the plasma membrane to other organelles (McMahon & Boucrot, 2011; Takamori *et al*, 2006). Specifically in neurons the synaptic delivery of proteins and lipids that are produced in the ER-Golgi complex, requires CME. Neurons also extensively use CME to recycle synaptic vesicles (SV) to maintain chemical communication (Heuser & Reese, 1973; Blondeau *et al*, 2004; Südhof, 2004; Granseth *et al*, 2006). Recent human genetics identified proteins mutated in- or associated with conferring risk to-neurodegenerative diseases. This is particularly relevant for Parkinson’s disease (PD) where mutations in the key CME-regulators Auxilin (*DNAJC6*, PARK19), Synaptojanin-1 (*SYNJ1*, PARK20) and EndophilinA-1 (*SH3GL2*) were identified, and also other accessory regulatory proteins to CME like LRRK2, RME-8, GAK (Auxilin-II) are implicated in this disease (Quadri *et al*, 2013; Krebs *et al*, 2014; Köroĝlu *et al*, 2013; Edvardson *et al*, 2012; Billingsley *et al*, 2018; Blauwendraat *et al*, 2020). However, how mutations in these proteins cause neuronal problems that are relevant to Parkinsonism, and whether there are functional relations and interactions between key “CME genes” mutated in PD, remains poorly understood.

Auxilin and Synj1 act in related steps of CME. During endocytosis, nascent vesicles can form because coat proteins like clathrin and adaptors help to shape the membrane at the Golgi complex or at the synaptic plasma membrane. Auxilin is a co-chaperone to Hsc-70 and is involved in rearranging the clathrin coat to remove it following vesicle formation (Fig 1A) (Jiang et al, 1997; Lee et al, 2006; He et al, 2019; Saheki & De Camilli, 2012). Synj1 is a phosphoinositide (PI) phosphatase that dephosphorylates vesicle membrane PIs causing the dissociation of proteins, like adaptors, that link the clathrin-lattice to the membrane bilayer (Saheki & De Camilli, 2012; McPherson et al, 1996; Verstreken et al, 2003; Woscholski et al, 1997) (Fig 1A). Interestingly, Auxilin also contains a motif to bind mono-PIs in its PTEN domain (Fig 1B). This suggests that Synj1-activity could promote Auxilin-recruitment to the membrane in the process of CME (Fig 1A).

**Figure 1.**
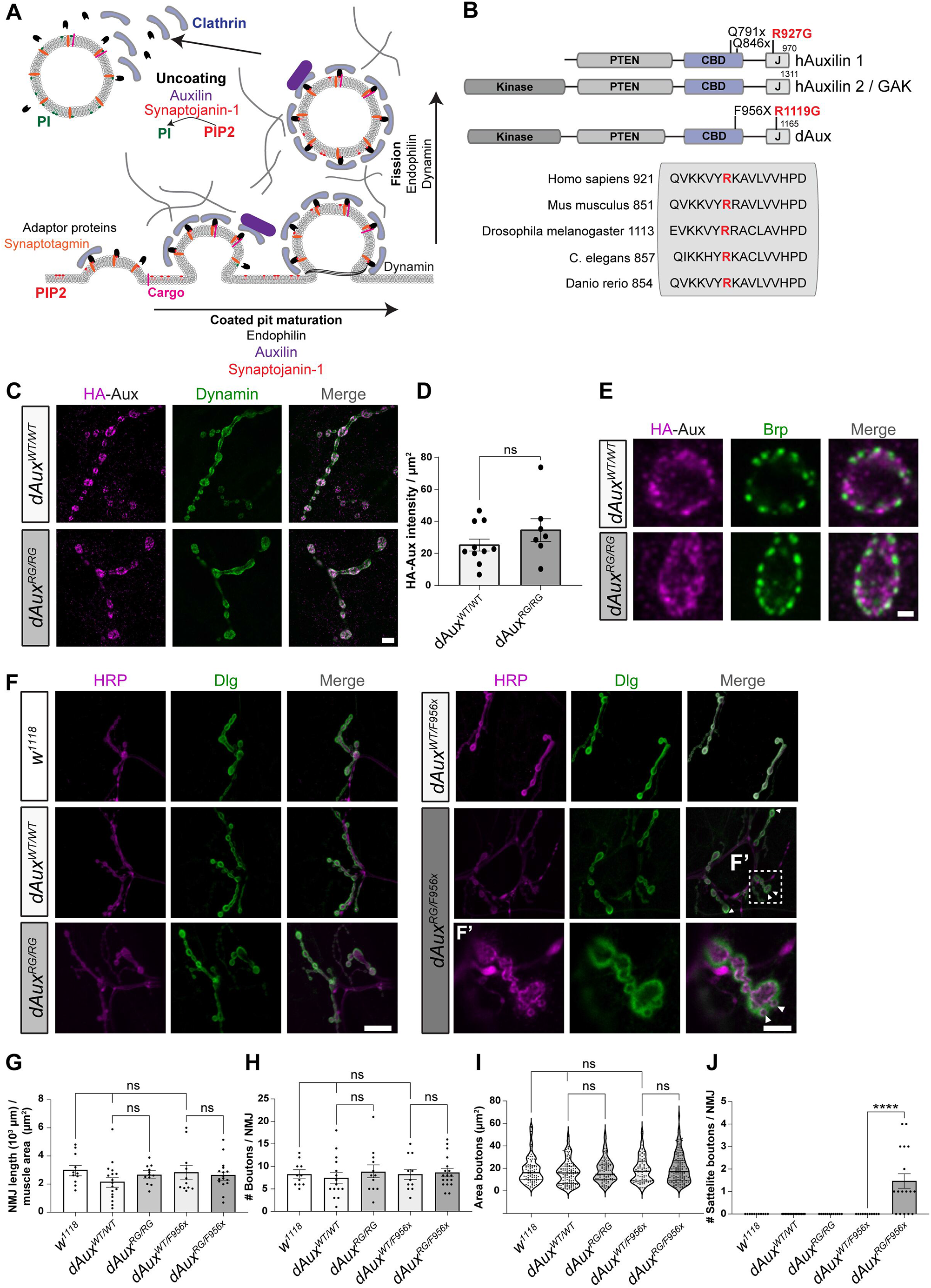
Auxilin protein level, localization and synaptic morphology are normal in pathogenic *dAux* mutant larvae. A. Schematic of clathrin-mediated-vesicle formation (here: endocytosis) (CME) indicating key proteins and lipids; conversion of phosphoinositides, phosphatidylinositol 4,5-bisphosphate [PI(4,5)P_2_] and phosphatidylinositol 3,4-bisphosphate [PI(3,4)P_2_] (PIP2). B. Domain structure of mammalian and *Drosophila* Auxilin and pathogenic mutations mentioned in this work. The arginine in human Auxilin that is mutated in PD, at position 927 (red) is conserved across species and homologous to *Drosophila* R1119. C-D. HA-tagged dAux^WT^ and dAux^RG^ proteins localize to the nerve terminals of the larval NMJs where they show overlap with Dynamin (Newmyer et al, 2003). (C) Representative images of NMJs labelled with anti-HA (magenta, dAux) and anti-Dynamin (green) and (D) quantification of the mean HA-Aux intensity per µm^2^. n≥8 larvae (of 3 independent crosses). Bars show the mean ± SEM, points show individual values. *t*-test, ns: not significant. Scale bar: 2.5 µm. E. Confocal images of L3 larval NMJs of *dAux*^*WT/WT*^ and *dAux*^*RG/RG*^ labelled with anti-Brp and anti-HA (magenta, dAux). Scale bar: 1 µm. F-J. Confocal images of NMJs of *w*^*1118*^, *dAux*^*WT/WT*^, *dAux*^*RG/RG*^, *dAux*^*WT/F956x*^ and *dAux*^*RG/F956x*^ larvae labelled with anti-Hrp and anti-Dlg (F) and quantification (G-J). (F’) Enlargement of the indicated panel, white arrow: satellite boutons. Quantification of NMJ length per muscle area (G), bouton number per NMJ (H), size of individual boutons (I) and total number of satellite boutons per NMJ (J). n=3 larvae (of 3 independent crosses). Dots represent the result of single NMJs (G, H & J) or area of single boutons (I). Bars show the group mean ± SEM, points show individual values. One-way ANOVA, with Dunnet’s multiple comparison test. ns: not significant, *** p < 0.001. Scale bars left 20 µm; right 5 µm.

While these functional studies relied on severe loss-of-function or null mutants that cause lethality (Hagedorn *et al*, 2006; Kandachar *et al*, 2008; Song *et al*, 2017; Verstreken *et al*, 2003; Cremona *et al*, 1999; Yim *et al*, 2010), pathogenic mutations in *DNAJC6* and in *SYNJ1* cause only partial inactivation of protein function (Cao *et al*, 2017; Roosen *et al*, 2021). Interestingly, in patients, these mutations result in a very similar spectrum of phenotypes that include “typical Parkinson defects” such as bradykinesia, resting tremor, rigidity and postural instability, as well as additional clinical problems including developmental delay and seizures (Edvardson *et al*, 2012; Olgiati *et al*, 2016; Köroĝlu *et al*, 2013; Olgiati *et al*, 2014; Kirola *et al*, 2016; Taghavi *et al*, 2018; Romdhan *et al*, 2018; Quadri *et al*, 2013; Krebs *et al*, 2014). However, the consequences these mutations have on synaptic integrity and neuronal survival during ageing, and whether Auxilin and Synj1 can compensate for one another in the context of disease remains enigmatic.

Creating new vesicles through CME involves topological membrane changes that highly depend on fine-tuned interactions between specific lipids and vesicle shaping- and budding-proteins. Recent observations recognize lipid disturbances in PD, including that there are changes in ceramide, sphingolipids, fatty acids, cholesterol and neutral lipids in *post mortem* brains of PD patients (Brekk *et al*, 2020; Fanning *et al*, 2020; Fais *et al*, 2021). Whether these changes are primary defects or “end-stage” and thus confounded by the long disease process, is unclear. It is also not known what the functional consequences of such changes are on cellular and organellar function. However, it is conceivable that lipid alterations affect vesicle trafficking, thus connecting these to PD.

Here, we find that the PD pathogenic mutation knocked into *Auxilin/DNAJC6* causes *in vivo* lipid disturbances, neuronal dysfunction, and neurodegeneration. *Drosophila dAux*^*RG*^ mutants have lower levels of membrane lipids containing long-chain polyunsaturated fatty acids (LC-PUFA), including PI lipid species. Synj1, also mutated in PD, preferentially binds LC-PUFA PIs, and we show that neuronal overexpression of Synj1 fully rescues the *dAux*^*RG*^ caused neuronal dysfunction, including behavioral defects and neurodegeneration. This work shows that the role of two PD genes is connected through neuronal lipid metabolism and suggests an important role for lipids in PD pathogenesis.

## RESULTS

### The pathogenic Auxilin mutant is a weak hypomorphic allele

Blunt loss of Auxilin function causes early lethality, but partial loss-of-function causes neurological problems (Edvardson *et al*, 2012; Olgiati *et al*, 2016; Köroĝlu *et al*, 2013; Hagedorn *et al*, 2006; Kandachar *et al*, 2008; Song *et al*, 2017; Yim *et al*, 2010). Among several pathogenic mutations that have been isolated in *DNAJC6*, the R927G mutation in the J-domain causes the most specific Parkinson’s disease problems (Olgiati *et al*, 2016) and the mutation is contained within the His-Pro-Asp (HPD) motif of the J-domain that is essential for Hsc70 recruitment (Fig 1B). To model *DNAJC6/Aux*-induced disease, we created new *Drosophila* knock-in animals that express the pathogenic *Aux* mutation under endogenous promotor control. We first replaced the entire *dAux* gene by an AttP flanked *white*^*+*^ gene and then used ΦC31 to substitute the *white*^*+*^ marker by the entire *dAux*^*WT*^ gene or by the mutant *dAux*^*R1119G*^ gene (hereafter referred as *dAux*^*RG*^); both genes are 5’-HA-tagged (R1119G is homologous to the human pathogenic R927G mutation (Fig 1B)).

To determine the impact of the R1119G mutation on synaptic function, we resorted to the third instar neuromuscular junction (NMJ) that is frequently used as a model synapse to study neuronal cell biology. We first examined the subcellular localization of dAux in *dAux*^*WT/WT*^ and *dAux*^*RG/RG*^ using anti-HA antibodies. dAux^WT^ and dAux^RG^ both localize to NMJ nerve terminals (Fig 1C & D), where they partially overlap with Dynamin, a known binding partner of Auxilin that acts in endocytosis (Fig 1A) (Newmyer et al, 2003). We also find dAux^WT^ and dAux^RG^ to be both enriched in the regions adjacent to active zones labelled by anti-Bruchpilot (Fig 1E), that are sites of synaptic vesicle endocytosis (Wagh *et al*, 2006). This indicates that dAux localizes to synapses, similar to its mammalian counterpart (Ahle & Ungewickell, 1990; Roosen *et al*, 2021) and the R1119G mutation does not impair protein stability or localization at peri-active zones.

The appearance of supernumerary “satellite boutons” on Type 1b boutons is often seen in endocytic and membrane recycling mutants (Collins & DiAntonio, 2007; Khodosh *et al*, 2006; Koh *et al*, 2004; Dickman *et al*, 2006; Budnik *et al*, 1990; Verstreken *et al*, 2003). To test if *dAux* mutants have morphological defects at their NMJs, we labelled wild type controls (*w*^*1118*^), *dAux*^*WT/WT*^ and *dAux*^*RG/RG*^ with anti-HRP and anti-DLG that mark pre- and post-synaptic sites of NMJ boutons (Fig 1F). Similar to *w*^*1118*^ and *dAux*^*WT/WT*^, satellite boutons do not appear in *dAux*^*RG/RG*^. Likewise, other morphological features, including the total number of boutons, bouton diameter, NMJ length or muscle surface area and length are not significantly different between these genotypes (Fig 1F-J). Hence, at the level of our analysis, homozygous mutant *dAux*^*RG/RG*^ have normal NMJ morphology and do not suffer from a developmental delay.

To further investigate the nature of the R1119G mutation we performed a genetic analysis. We combined *dAux*^*RG*^ and *dAux*^*WT*^ with the severe hypomorphic *dAux*^*F956X*^ mutation (Kandachar *et al*, 2008). *dAux*^*F956X*^ harbors a stop codon after the clathrin binding domain, thus removing the J-domain (Fig 1B). Synaptic morphology of *dAux*^*WT/F956X*^ and *dAux*^*RG/F956X*^ is very similar (Fig 1F-I), except there are significantly more satellite boutons in *dAux*^*RG/F956X*^ (Fig 1F, F’ white arrows and J). These data are consistent with *dAux*^*RG*^ being a weak hypomorphic (loss-of-function) allele whose phenotypes are revealed when combined with more severe alleles.

### Pathogenic Auxilin mutants harbor irregular sized synaptic vesicles

To test if the Auxilin pathogenic mutant affects synaptic vesicle (SV) endocytosis, we quantified the uptake of a fluorescent lipophilic dye, FM1-43, that is internalized by endocytosis into SVs upon stimulation (10 min, 90mM KCl, 1.5mM CaCl_2_) (Betz & Bewick, 1992). Under these conditions, no difference in dye uptake is observed between *w*^*1118*^ controls, *dAux*^*WT/WT*^, *dAux*^*RG/RG*^ *dAux*^*WT/F956X*^ and *dAux*^*RG/F956X*^ mutants (Fig EV1A & B).

To reveal the ultrastructure of the Auxilin mutant synapses we used transmission electron microscopy. We activated the vesicle cycle and thus endocytosis by first stimulating the NMJs before processing and imaging (Fig EV1C). We then implemented a machine learning approach to quantify the data in an unbiased manner (see methods). Ultrastructural features such as pre-synaptic active zones and T-bars, and mitochondrial surface area (Fig EV1C) and cristae structure are not different between the analyzed genotypes. Similarly, we do not find differences in SV number (<80 nm) or cisternae number (>80 nm) (Fig EV1D-F) that are both parameters often found affected in endocytic mutants (Kadota & Kadota, 1982; Koenig & Ikeda, 1996; Kasprowicz *et al*, 2008; Glyvuk *et al*, 2010; Meunier *et al*, 2010; Vanhauwaert *et al*, 2017). Next, we quantified the relative frequency of SV (<80 nm) diameter in bins of 5nm. These data reveal a more heterogeneous vesicle population, including a shift towards smaller SVs in *dAux*^*RG/RG*^ and in *dAux*^*RG/F956X*^ mutants, compared to *w*^*1118*^, *dAux*^*WT/WT*^, and *dAux*^*WT/F956X*^ controls (Fig EV1D & G). While the FM 1-43 dye uptake experiment suggests that the total amount of membrane uptake during stimulation is not affected by the pathogenic Auxilin mutant, the irregularly sized vesicle population that forms, is consistent with an impairment of efficient clathrin coat re-arrangements during vesicle formation.

### The Auxilin R1119G mutation causes severe behavioral dysfunction

The larval NMJ is a very robust synapse that may not reveal subtle problems arising during ageing. We therefore turned to adult flies and performed a genetic complementation analysis using *dAux*^*WT*^, *dAux*^*RG*^, *dAux*^*F956X*^ and *dAux*^*-*^ null mutant flies. While *dAux*^*–/–*^, *dAux*^*F956x/–*^ and *dAux*^*RG/–*^ is lethal, a few *dAux*^*RG/F956X*^ mutant animals escape lethality, but all these flies die within 25-days-post-eclosion. Conversely, *dAux*^*WT/-*^, *dAux*^*WT/F956X*^, *dAux*^*WT/RG*^ *and dAux*^*RG/RG*^flies are viable (Fig 2A). Hence, the data suggests the following allelic series: *dAux*^*WT*^ > *dAux*^*RG*^> *dAux*^*F956X*^ > *dAux*^*-*^.

**Figure 2.**
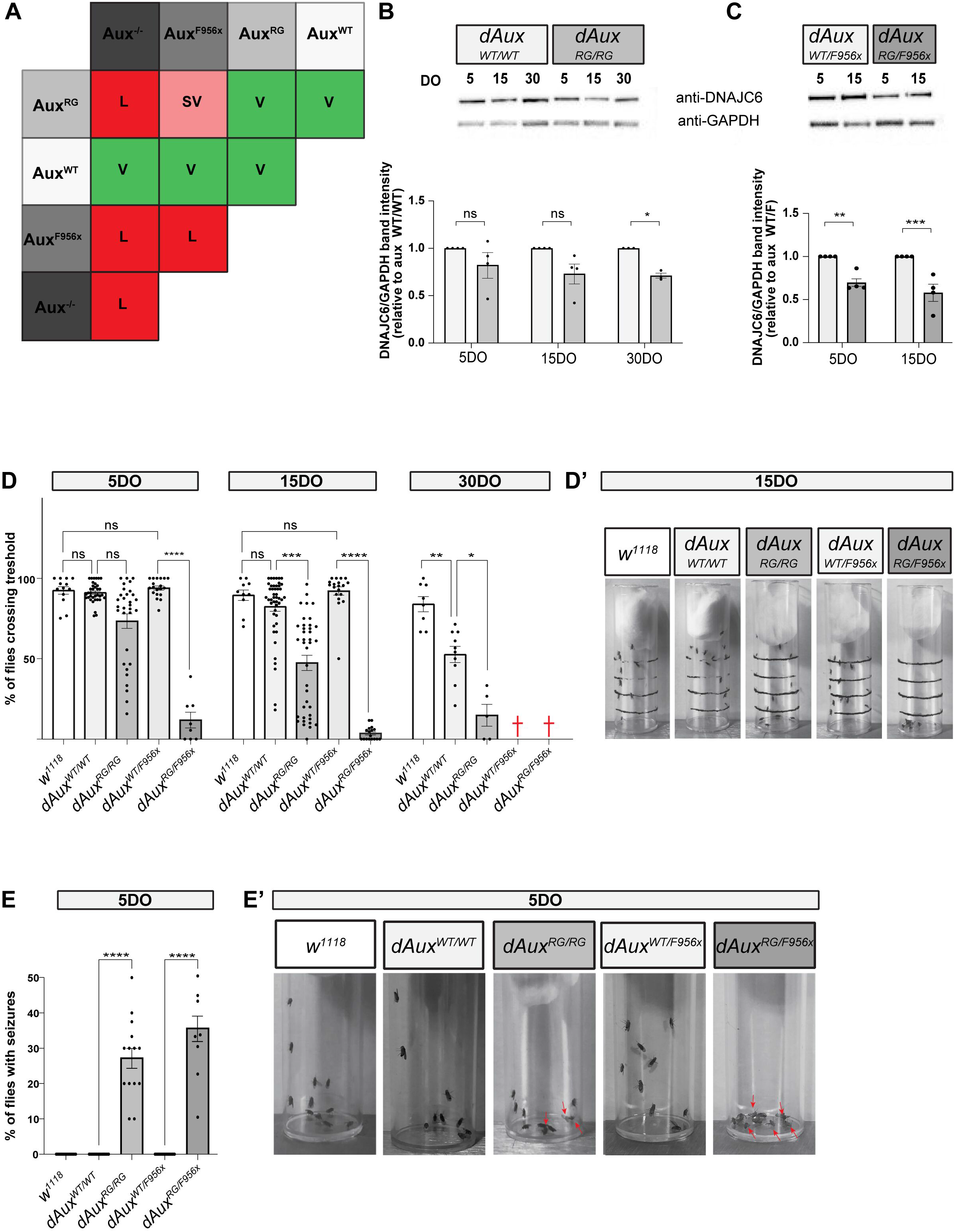
The pathogenic *Auxilin* mutants develop age-dependent neurological abnormalities. A. Complementation table indicating viability/lethality of the indicated genotypes. L: Lethal, SV: semi-viable and V: viable. B-C. Images of Western blots from brain lysates prepared from 5, 15 and 30DO flies of indicated genotypes labelled with anti-DNAJC6/Auxilin and anti-GAPDH (loading control) (B) and quantification of Auxilin protein levels (C). Values are relative to GAPDH and are expressed as a fraction of protein levels in *dAux*^*WT/WT*^. Bars show the mean ± SEM, points show individual values and n≥3. Two-way ANOVA, with Bonferroni’s post hoc test. *p<0.05,**p<0.01 and ***p<0.001. D. Quantification of the negative geotaxis assay of 5DO, 15DO or 30DO flies of the indicated genotypes (D) and representative images of 15DO flies after 30s (D’). Graphs represent the fraction of flies crossing the 4cm mark. Note that most *dAux*^*WT/F956X*^ and all *dAux*^*RG/F956X*^ flies die within 25-days-post-eclosion (red crosses). Bars show mean ± SEM. Points represent the mean of three trials each. One-way ANOVA, Dunn’s multiple comparison test. ns: not significant, *p<0.05, **p<0.01, ***p<0.001 and ****p<0.0001. E. Quantification of seizures following vortexing in 5DO flies of indicated genotypes (E) and representative images of flies after vortex-stimulation (E’). Graphs represent the fraction of flies that were unable to stand 10 s after stimulation. Bars show mean ± SEM. Points represent single trials with 10 flies each. Red arrows: flies that are not able to stand. One-way ANOVA, Dunnett’s multiple comparison test. ns: not significant, ****p<0.0001.

To determine if the pathogenic mutation affects Auxilin protein level over time, we conducted Western blotting of 5, 15 and 30 day-old (DO) adult head tissue of *dAux*^*WTWT*^, *dAux*^*RG/RG*^, *dAux*^*WT/F956X*^ and *dAux*^*RG/F956X*^ (the latter two only at 5 and 15 DO). Interestingly, a decrease of Auxilin protein levels is observed over time, when compared to controls (Fig 2B & C). Furthermore, the drop in protein levels is more pronounced when the R1119G and F956X mutations are combined compared to homozygous *dAux*^*RG/RG*^ (Fig 2B & C). This further confirms that R1119G is a weak hypomorpic mutant and that it leads to a small age-dependent decline in protein abundance.

We next asked if the mutant *dAux* flies display progressive neurological problems, including motor defects and seizure-like behavior. In negative geotaxis assays, 5DO *dAux*^*RG/RG*^ flies can climb past a 4 cm mark within a 30 s window, however *dAux*^*RG/F956X*^ flies fail to do so (Fig 2D & D’). Furthermore, this is progressive, and the ability to reach the 4 cm mark declines with age for both *dAux*^*RG/RG*^ and *dAux*^*RG/F956X*^ (Fig 2D & D’). *DNAJC6/Auxilin* PD patients also frequently suffer from seizures. We therefore assessed this phenotype in 5DO flies by triggered by 10 s of vortexing. This causes strong seizures in both *dAux*^*RG/RG*^ and *dAux*^*RG/F956X*^ flies, but not in the controls (Fig 2E & E’). These behavioral tests indicate that the pathological *DNAJC6/Auxilin* mutation causes neurological defects that resemble those observed in patients, including motor impairments and seizures.

### Auxilin R1119G causes neurodegeneration

To determine the integrity of neuronal function in adult flies, we performed electroretinogram (ERG) recordings. ERGs assess neuronal function and integrity of photoreceptors (the depolarization phase “DEP”) and synaptic transmission in the visual system (“ON” and “OFF” transients – red arrows) upon exposure to a short light pulse (1 s) (Verstreken *et al*, 2003) (Fig 3A). 5DO *dAux*^*RG/RG*^ flies show a stereotypical response, similar to *dAux*^*WT/WT*^ (Fig 3A). However, 5DO *dAux*^*RG/F956X*^ flies show a lower DEP amplitude and smaller ON and OFF transients compared to controls (Fig 3A). This phenotype worsens progressively, and 15 and 30DO animals show gradually smaller ON and OFF transient amplitudes (Fig 3A). Finally, this defect is completely rescued by transgenic expression of wild-type *dAux* in neurons using *nSyb-Gal4* (Fig EV2A). These data indicate that the pathogenic mutation in Auxilin causes a cell autonomous and progressive loss of neuronal integrity in mutant adult flies.

**Figure 3.**
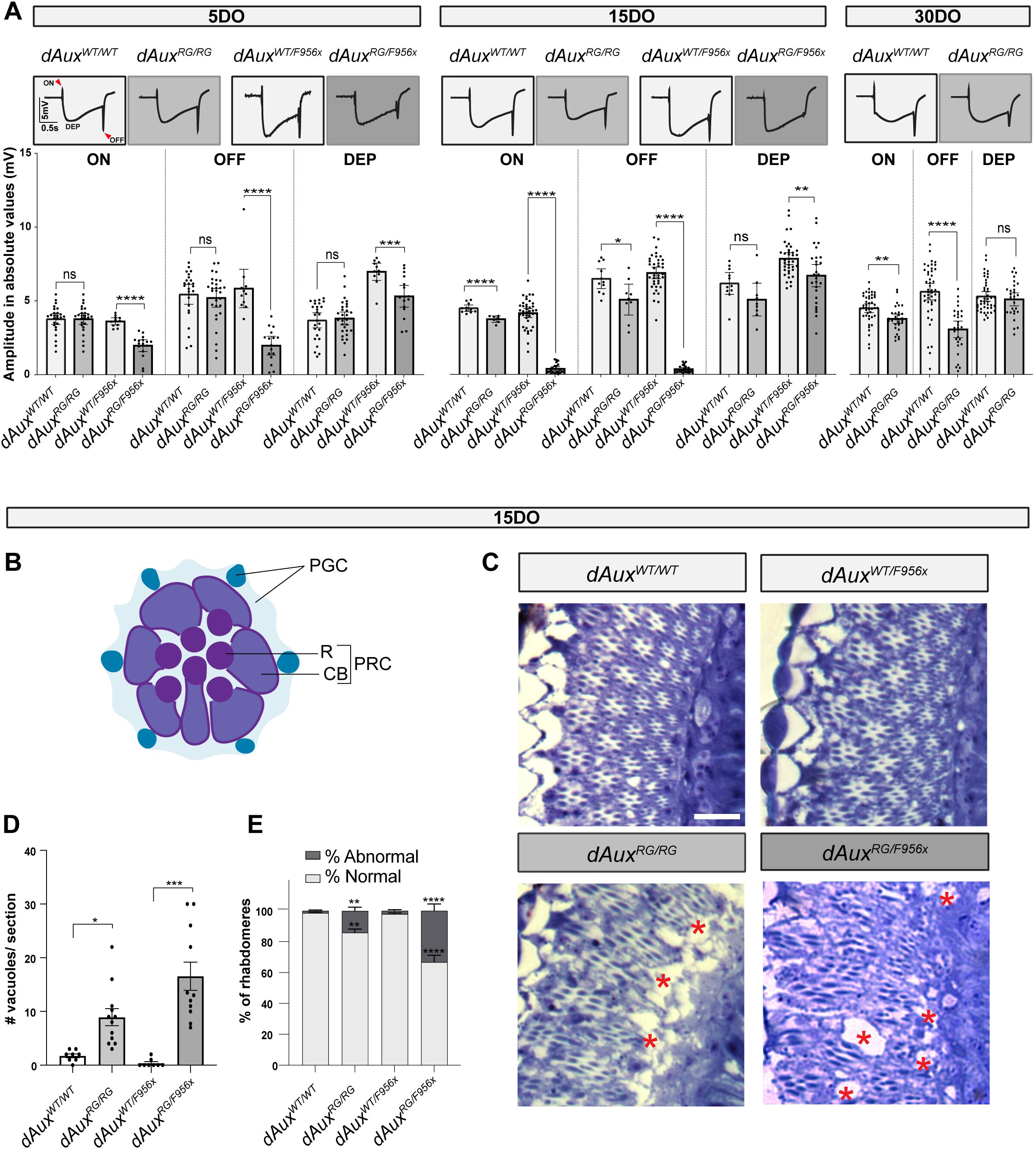
Pathogenic *Auxilin* mutants show neuronal function defects and neurodegeneration. A. Average ERG traces of 5, 15 and 30DO flies of indicated genotypes and quantification on ON, OFF and depolarization amplitude. Arrowheads: “ON” and “OFF” transients. Bar graphs represent absolute mean ± SEM values (mV). n≥10 flies per genotype, points represent values obtained from individual flies. One-way ANOVA, Dunnett’s multiple comparison test. ns: not significant, ** p < 0.01, *** p < 0.001 and **** p < 0.0001. B. Diagram of a tangential section of the adult *Drosophila* retina, with pigmented glia cells (PGC) in blue and photoreceptor cells (PRC) in purple. R: rhabdomere; CB, cell body. C-E. Representative brightfield images of toluidine blue stained sections of 15DO *dAux*^*WT/WT*^, *dAux*^*RG/RG*^, *dAux*^*WT/F956X*^ and *dAux*^*RG/F956X*^ fly eyes (C; asterisks indicate neurodegeneration) and quantification of the degenerated area (dubbed vacuoles in the literature/number of vacuoles counted) per standardized section (D) and the amount of normal *versus* abnormal rhabdomeres (methods) as a fraction of the total number of rhabdomeres (E). The graph in D shows the mean ± SEM of n≥10 flies per genotype from independent crosses and points are individual values. Kruskal-Wallis test, Dunnett’s multiple comparison test. * p < 0.05 and *** p < 0.001; the graph in E shows mean ± SEM of n≥5 flies per genotype from independent crosses. Two-way ANOVA with Tukey’s multiple comparison test. ** p < 0.01 and **** p < 0.0001. Scale bar in C is 5 µm.

ON and OFF transient defects in ERG recordings arise from defects in synaptic transmission (Verstreken *et al*, 2003; Uytterhoeven *et al*, 2015), consistent with the known role of Auxilin in ensuring an efficient synaptic vesicle cycle (He *et al*, 2019; Lee *et al*, 2006; Massol *et al*, 2006). The smaller depolarisation in ERG recordings suggests the *dAux* mutants cause neurodegeneration. We therefore prepared serial histological sections of 15DO animals and stained them with toluidine blue. Quantification revealed significantly more degeneration (“vacuoles”) in the optic lobes of *dAux*^*RG/RG*^ and *dAux*^*RG/F956X*^ flies compared to the controls (Fig 3B-D) (Sunderhaus & Kretzschmar, 2016). Furthermore, we also observe a loss of structural integrity in the compound eye. We find that 15DO *dAux*^*RG/RG*^ and *dAux*^*RG/F956X*^ animals have an increased number of abnormal ommatidia with fewer than 7 normally positioned rhabdomeres in a section (Fig 3C & E). Taken together, the pathogenic mutation in *DNAJC6/Auxilin* causes a loss of neuronal integrity and neurodegeneration.

### Pathogenic Auxilin mutations affect lipids that are critical to Synaptojanin function

*DNAJC6/Auxilin* acts to remove clathrin coats at endocytic membranes. To investigate if the pathogenic mutation in *DNAJC6/Auxilin* is associated with membrane lipid changes, we performed shotgun lipidomics of 15DO wild type and mutant fly heads and examined membrane glycerol- and glycerophospholipids (GL and GPL). The major membrane GPLs we find in wild-type adult fly head are phosphatidylethanolamine (PE, ∼50%), phosphatidylcholine (PC, ∼29%), phosphatidylserine (PS, ∼12%) and phosphatidylinositol (PI, ∼4%) (Fig 4A). The GPLs phosphatidic acid (PA) and phosphatidylglycerol (PG) and the GL diacylglycerol (DAG) are present in lower amounts (Fig 4A). We find a similar relative abundance of GL and GPL in *dAux*^*RG/RG*^ and *dAux*^*RG/F956X*^ flies, except for the minor lipid species, PA, that is reduced in *dAux*^*RG/F956X*^ mutants (Fig 4B). This indicates that the composition of the major membrane phospholipids is not strongly affected by the pathogenic Aux mutation.

**Figure 4.**
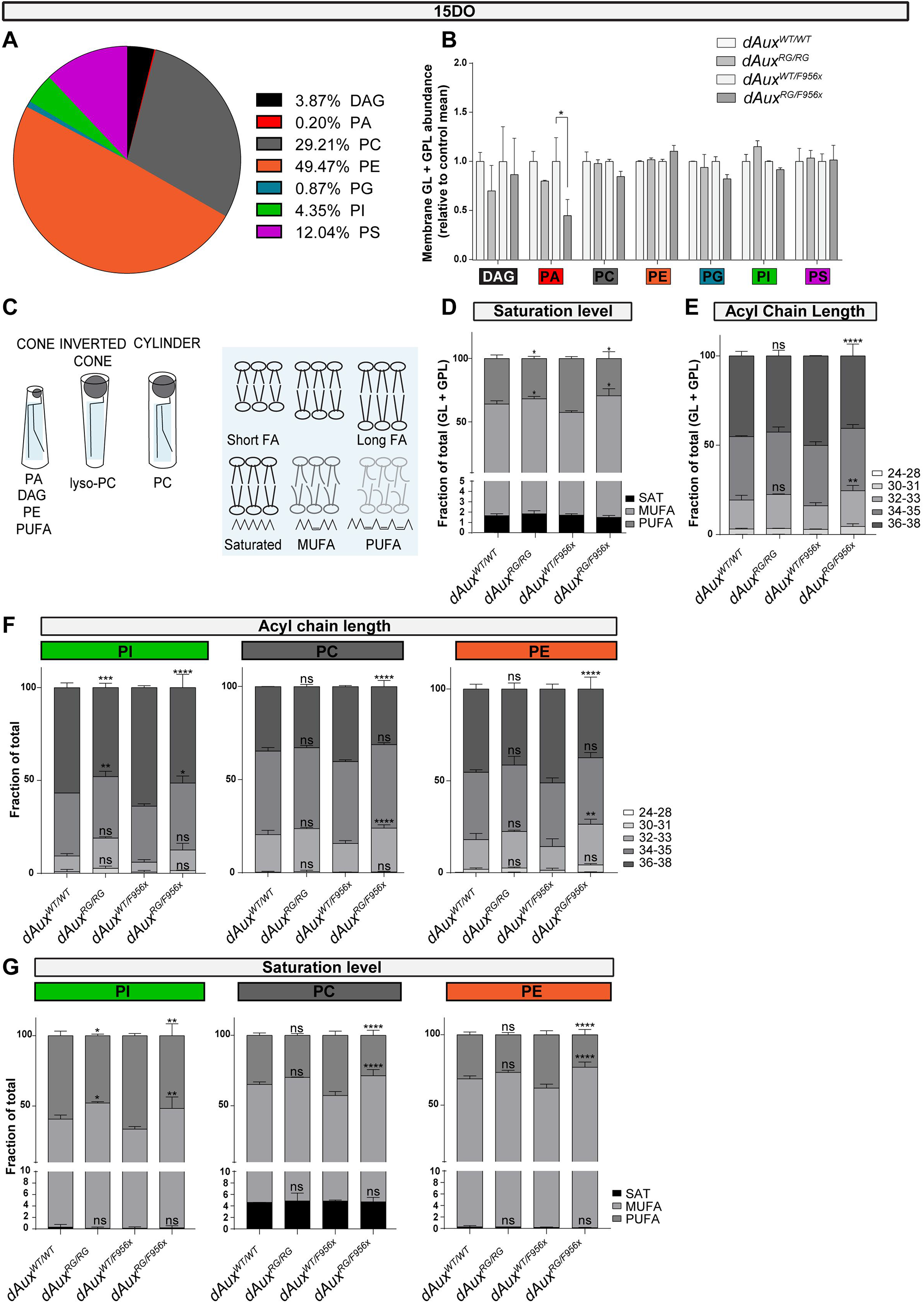
Membrane lipids containing Long-Chain Polyunsaturated Fatty Acids, including phosphoinositides species are decreased in pathogenic *Auxilin* fly heads. A. Pie diagram of the relative abundance of individual GL and GPL classes in 15DO wild-type fly heads detected by mass spectrometry (MS). n = 2 analyses and one individual analysis is performed on 15 fly heads collected from 3 independent crosses. B. The relative abundance of individual GL and GPL classes in 15DO fly heads of indicated genotypes relative to *dAux*^*WT/WT*^ and as detected by mass spectrometry (MS). Bars show mean ± SEM of n = 2 analyses. One individual analysis is performed on 15 fly heads collected from 3 independent crosses. Two-way ANOVA, Tukey’s multiple comparison test. *p<0.05. C. Schematic of the geometry of membrane lipids, showing the effect of polar headgroups and fatty acyl side chains. Membrane thickness, fluidity and deformity is influenced by the saturation level and length of the fatty acyl chain in concert with the presence of cholesterol (Antonny *et al*, 2015). D-E. Saturation level (D) and length (E) of fatty acyl chains of all membrane lipids are presented as fraction of total GL+GPL. Bars show mean ± SEM. = 2 and one individual analysis is performed on 15 fly heads. MUFA, mono-unsaturated; PUFA, poly-unsaturated. Two-way ANOVA, Tukey’s multiple comparison test. ns: not significant, *p<0.05, **p<0.01 and ****p<0.0001. F-G. Length (F) and saturation level (G) of fatty acyl chains of PI, PE, PC are presented as faction of total GPL. Note the lower levels of LC-PUFAs including PI LC-PUFAs in *dAux* mutants. Bars show mean ± SEM. n = 2 analyses and one individual analysis is performed on 15 fly heads collected from 3 independent crosses. Two-way ANOVA, Tukey’s multiple comparison test. ns: not significant, * p < 0.05, ** p < 0.01, *** p < 0.001 and **** p < 0.0001.

Membrane properties such as thickness, fluidity and curvature that affect synaptic function are dictated by lipid shape (eg cone- and inverted cone shaped lipids - Fig 4C; Antonny et al, 2015), but also by the fatty acid chain saturation level and length (Fig 4C; Antonny et al, 2015). Furthermore, membrane lipids containing long-chain polyunsaturated fatty acids (LC-PUFAs) that make up ∼30% of the brain lipids and are also important to maintain a normal SV pool (Sastry, 1985; Qi *et al*, 2002; Marza & Lesa, 2006; Pinot *et al*, 2014; Takamori *et al*, 2006; Yang *et al*, 2012). In 15DO *dAux*^*RG/RG*^ and *dAux*^*RG/F956X*^ mutant fly heads we find an increase of mono-unsaturated fatty acids (MUFA) and a significant decrease in poly-unsaturated fatty acids (PUFA) in all GLs and GPLs, compared to controls (Fig 4D).

Next, we assessed fatty acid chain length and find that membrane lipids (GL + GPL) are shorter in *dAux*^*RG/F956X*^ mutants compared to controls (Fig 4E). The lack of a significant change in fatty acid chain length in *dAux*^*RG/RG*^ could be masked since we analysed GL and GPL in bulk. We thus also investigated saturation level and chain length per lipid class. The most prominent change is a significant decrease in LC-PUFA PI species in both *dAux*^*RG/RG*^ and *dAux*^*RG/F956X*^ mutants compared to controls (Fig 4F & G). More importantly, changes in PI 36:4 and PI 32:1 levels are detected in both *dAux*^*RG/RG*^ and *dAux*^*RG/F956X*^ mutants, but also other lipid species show changes (Fig EV3A-C). This indicates that the pathogenic mutation in *DNAJC6/Auxilin* induces evident changes in lipid abundance and structure, including in lipid species that are required at synapses (LC-PUFA PI species).

### Neuronal Synaptojanin expression rescues pathogenic *DNAJC6/Auxilin* mutant defects

Membrane lipids containing LC-PUFA and PI lipid species play a prominent role in neurons and at synapses (Cremona *et al*, 1999; Khuong *et al*, 2013); several endocytic proteins interact with these lipids, including Synj1. Lipids containing LC-PUFA are involved in localizing Synj1 to sites of neurotransmitter release (Marza *et al*, 2008; Schmid *et al*, 2004) and Synj1 also binds to and dephosphorylates PI (Posor *et al*, 2015; Salim *et al*, 1996; Marza *et al*, 2008; Schmid *et al*, 2004). Given that patients with mutations in *SYNJ1* show clinical phenotypes very similar to patients with mutations in *DNAJC6* (Kirola et al, 2016; Taghavi et al, 2018; Romdhan et al, 2018; Quadri et al, 2013; Krebs et al, 2014), we asked if the pathogenic *dAux*^*RG*^ mutation genetically interacts with *Synj*. We created *dAux*^*RG/RG*^ and *dAux*^*RG/F956X*^ animals that overexpress Synj in neurons (using *nSyb-Gal4*). As a control we also created *dAux*^*RG/RG*^ and *dAux*^*RG/F956X*^ animals that overexpress Aux in neurons. We find that either expression of Aux or of Synj in *dAux* mutants are each able to rescue the negative geotaxis defects and prevent seizures in 5DO mutant flies (Fig 5A & B). Similarly, also the neurodegeneration defects, measure by ERG and histology, of 15DO *dAux*^*RG/RG*^ and *dAux*^*RG/F956X*^ mutants are rescued by neuronal expression of Synj (Fig 5C-E and Fig EV4A-D). In contrast, overexpression of clathrin heavy chain, another Auxilin-interacting protein that is not mutated in PD, does not rescue the defects in *dAux*^*RG/RG*^ and *dAux*^*RG/F956X*^ mutants (Fig 5A-E and Fig EV4A-D). These results indicate that Synj can compensate for the behavioral, functional and neurodegeneration defects induced by the pathogenic *Auxilin/DNAJC6* mutations.

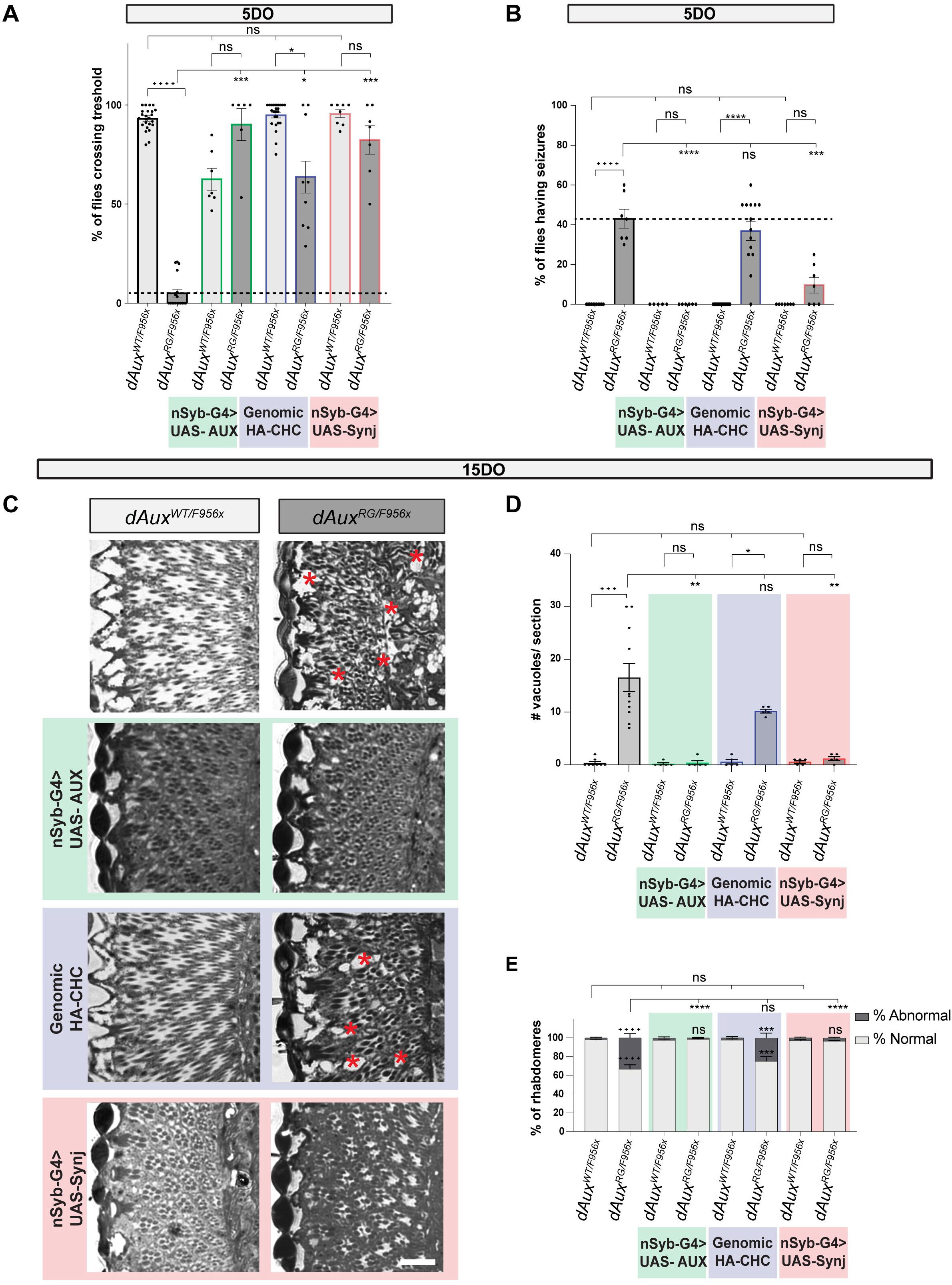

## DISCUSSION

In this work, we find that overexpression of the lipid phosphatase Synj fully rescues the dysfunction induced by a pathogenic mutation in *Auxilin/DNAJC6*. Auxilin regulates vesicle endocytosis/budding, and we uncovered in *dAux* mutants specific lipid changes, including PIs and LC-PUFAs, that are required for the function of Synj. These findings suggest important roles for lipid metabolism in the pathogenicity of PD and, like the discovery that expression of the PD gene *parkin* rescues aspects of *pink1* deficiency (Clark *et al*, 2006; Park *et al*, 2006; Yang *et al*, 2006), our work now uncovers a novel functional relation between two other Parkinsonism genes that both cause a similar early onset disease and clinical manifestation (Edvardson *et al*, 2012; Olgiati *et al*, 2014; Quadri *et al*, 2013; Olgiati *et al*, 2016; Köroĝlu *et al*, 2013; Kirola *et al*, 2016; Taghavi *et al*, 2018; Romdhan *et al*, 2018; Krebs *et al*, 2014).

There are several pathogenic mutations in *DNAJC6*; all share Parkinsonism phenotypes, but unlike the R927G (R1119G) mutation that we modelled here, other pathogenic mutations result in more severe additional neurological defects (Edvardson *et al*, 2012; Köroĝlu *et al*, 2013; Olgiati *et al*, 2016; Elsayed, 2016). Our data reveal that the R927G (R1119G) mutation is a weak loss-of-function mutation, in line with the relatively slow progression of the neurodegenerative features, as opposed to the very early onset neurological defects that are observed in the more severe loss-of-function mutations in *DNAJC6* (eg Q734X and Q789X; Köroĝlu et al, 2013; Elsayed, 2016).

A common feature across the *DNAJC6* mutations that cause Parkinsonism is that they affect the Auxilin J-domain (Fig 1B). The J-domain is a co-chaperone domain that functions with Hsc-70 and the best characterized function is its role in rearranging and dissolving the clathrin basket to support uncoating of newly formed vesicles (Jiang *et al*, 1997). In the weak hypomorphic and pathogenic R927G (R1119G) *Auxilin* mutant animals we did not find strong endocytic defects. However, we did observe smaller sized synaptic vesicles that could be the result of inefficient clathrin re-arrangements at the membrane. This may affect coated pit maturation, leading to defective vesicle constriction and the observed effects on SV size. It could equally affect other budding events in the cell, including the formation of Golgi-derived vesicles, where Auxilin is also known to act (Zhou *et al*, 2011; Ding *et al*, 2016).

Our work puts lipid changes central to synaptic and neuronal dysfunction in Parkinson’s disease. We report that the R927G (R1119G) pathogenic mutant harbors lower amounts of GL and GPL containing LC-PUFAs. The majority of lipid metabolic enzymes exert their function at the ER, and thus this is the main center of GLs and GPL generation. Lipids are then delivered to different locations including Golgi and synaptic membranes through lipid transfer proteins and transport vesicles (Hanada, 2018; Jacquemyn et al, 2017; Wong et al, 2017). In recent work on BioRxiv, Auxilin R927G knock in mice are suggested to harbor Golgi vesicle budding defects (Roosen *et al*, 2021), that thus would result in reduced lipid delivery, and explain why we detect changes in lipid abundance in *dAux*^*RG*^ mutants. Excitingly, we show that the upregulation of a single lipid phosphatase that is also mutated in Parkinson’s disease, Synaptojanin-1, rescues *Auxilin/DNAJC6*-induced neurodegeneration, pointing to a genetic interaction between these genes. This is also supported by observations that *Aux*^*KO*^ and *Synj*^*RQ*^ mice show similar dystrophic neurons in relevant nigrostriatal pathways and that *Synj1*^RQ^ mice have increased levels of Auxilin (Cao *et al*, 2017; Vidyadhara *et al*, 2022; Yim *et al*, 2010). Further data on BioRxiv, additionally suggests a synergistic relation between the two loss-of-function mutants: double mutants show more severe phenotypes; however, this work did not test if Synj expression rescues *Aux* mutant defects as we did here (Ng *et al*, 2022).

Our finding of specific lipid changes (lower levels of GL and GPL containing LC-PUFA, including lower levels of PI lipid species) in the pathogenic *DNAJC6/Auxilin* mutants may start to explain why we find that Synj expression strongly rescues pathogenic *dAux* mutants. PI metabolism is critical for numerous neuronal functions. They regulate the progression of endocytosis and vesicle budding (Posor et al, 2015; Salim et al, 1996; Marza et al, 2008; Schmid et al, 2004), the positioning of the exocytic machinery (Khuong *et al*, 2013), endosomal function (Wenk & De Camilli, 2004), autophagosomal function (Vanhauwaert *et al*, 2017), etc. Interestingly, we found decreased levels of PI LC-PUFA species. Phospholipids with LC-PUFA are enriched at the nerve terminals and are highly abundant in synaptic vesicles (Yang *et al*, 2012; Marszalek & Lodish, 2005; Takamori *et al*, 2006). They are known to facilitate membrane fission, including the function of the dynamin complex (Pinot *et al*, 2014). Importantly, Synj is known to bind preferentially LC-PUFA PI species to exert its functions (Marza *et al*, 2008; Schmid *et al*, 2004). Thus, we surmise that overexpression of Synj rebalances the system (possibly also at the Golgi) to support these activities. The model we propose is that pathogenic *Aux/DNAJC6* mutations affect lipid delivery that exacerbate SV recycling, and that increasing Synj1 levels draw more (mutant) Aux to the membrane, “fixing” dysfunction. Open questions for future work will be to understand how trafficking and lipid defects lead to degeneration, why some cells are more vulnerable to these defects than others, and whether similar lipid defects are also relevant in other (familial or idiopathic) forms of Parkinsonism.

## MATERIAL AND METHODS

### Fly Stocks, maintenance, and tissue collection

*Drosophila* stocks (table below) were maintained using standard protocols and fed a standard diet consisting of cornmeal, agar, yeast, sucrose, and dextrose. Experimental crosses were kept at 25°C. L3 larvae of desired genotypes were selected to perform immunostaining, FM-43 dye uptake assays and TEM. To conduct behavioural assays, Western blotting, ERGs, toluidine blue staining to assess neurodegeneration and lipidomics, adult flies of desired genotypes were selected after eclosion and aged in groups of 10-20 flies for 5, 15, or 30 days at 25°C. Flies were transferred to new vials every 3 days.

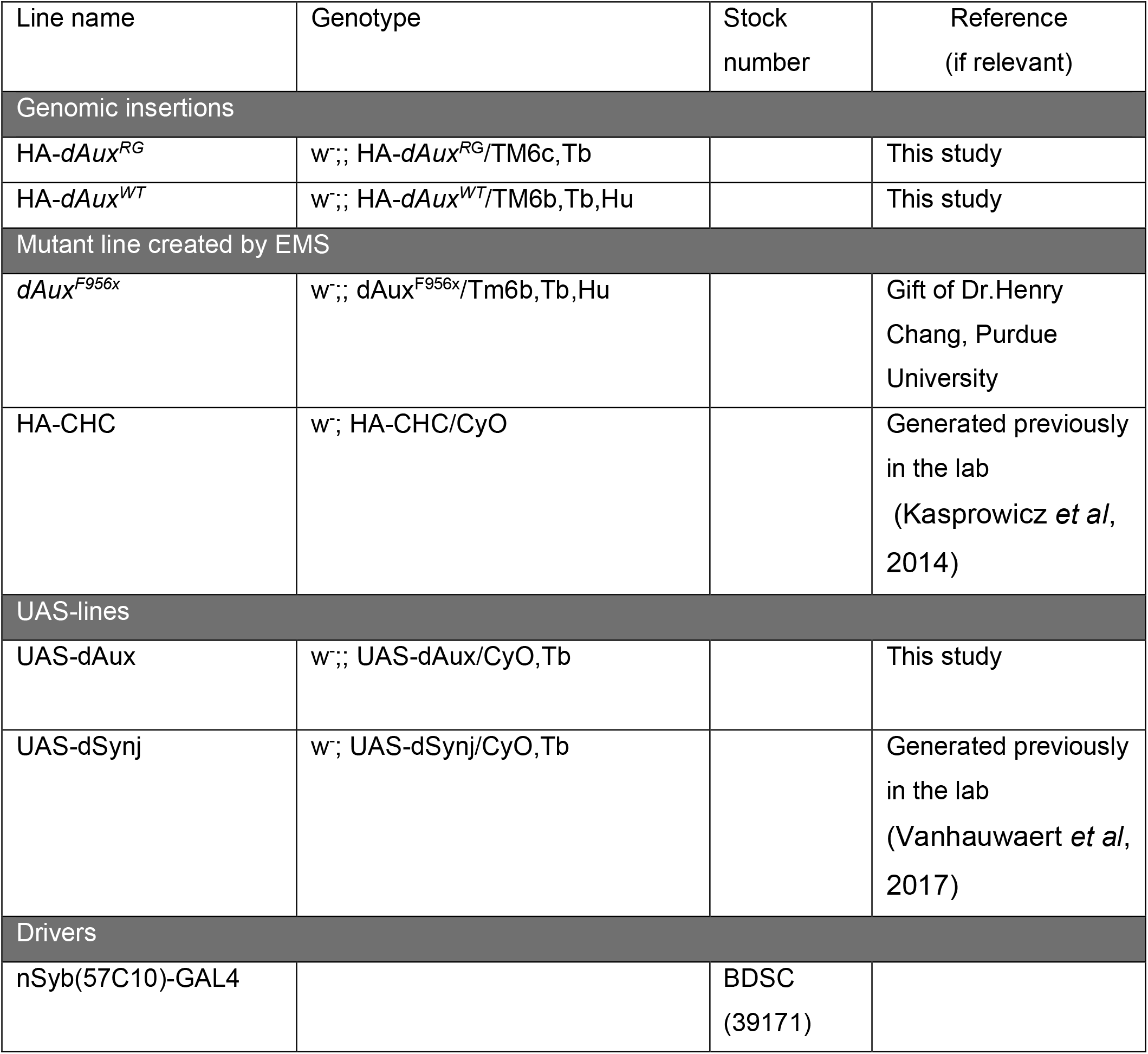

### New *Drosophila* Lines

Cloning was performed with the Gibson Assembly Master Mix (New England Biolabs). PCR products were produced with the Q5 High-Fidelity 2x Master Mix (New England Biolabs). Some fragments were ordered as gBlocks at IDT. All inserts were verified by sequencing. Microinjections were performed by BestGene Inc. (CA, USA). We used CRISPR-Cas9 to disrupt genomic dAux gene by injecting vas-cas9 (X chr) flies (Norrander *et al*, 1983) with pCFD3-dU6:3gRNA (Addgene #49410; (Port *et al*, 2014)) carrying guide RNA sequences and a pWhiteStar donor plasmid carrying an integrase mediated cassette exchange (IMCE) and a *white* gene. Knock-ins were created by injecting pBS-KS-attB1-2 (Addgene#61255; (Zhang *et al*, 2014)) containing HA-tagged *dAux*^*WT*^ or HA-tagged *dAux*^*RG*^ (the R1119G mutation) into recombinant vas-cas9:Phi31 flies. Transgenic fly lines were produced by injecting plasmids into VK37 for PhiC31 integrase-mediated site-specific transgenesis.

### HA-dAux^WT^ and HA-dAux^RG^

Unique gRNAs to disrupt the *dAux* gene were identified by flyCRISPR Target Finder Tool (https://flycrispr.org/) and predicted to introduce double-stranded breaks 249bp and 222bp before start and stop codon respectively. gRNAs were cloned into pCFD4:U6:1-gRNA U6:3-gRNA according to a previously established protocol (Port *et al*, 2014).

To generate a donor plasmid, DNA Assembly (NEB) was performed with four fragments: mini-*white* IMCE-cassette, left homology arm, right homology arm and pWhiteStar backbone (Choi *et al*, 2009). Left and right homology arms (1kb from gRNAs cutting side) of the *dAux* gene were ordered as gBlock at IDT. To avoid cutting the donor plasmid, the PAM sequences were mutated. The mini-*white* IMCE-cassette and vector backbone were generated by restriction digest on pWhite-STAR with AvrII and XhoI.

Next, a plasmid to exchange the mini-*white* IMCE-cassette was created by amplification of the entire *dAux* gene region from BAC CH322-22D05 (PACMANFLY, (Venken *et al*, 2009)) with an N-terminal primer containing the HA-tag and cloning this into our pBS-KS-attB1-2 using BbsI. The R1119G mutation in *dAux* was produced by performing Q5® Site-Directed Mutagenesis (NEB).

### UAS-dAux

The coding sequence of *dAux* (NM_164317) was amplified from a pOT2 vector (DGRC, SD05837) and cloned between the EcoRI and XhoI sites of pUAST-attB (Bischof *et al*, 2007) using the Gibson NEBuilder® Hifi DNA assembly kit. The coding region of pUAST plasmids was fully verified by Sanger sequencing before further use.

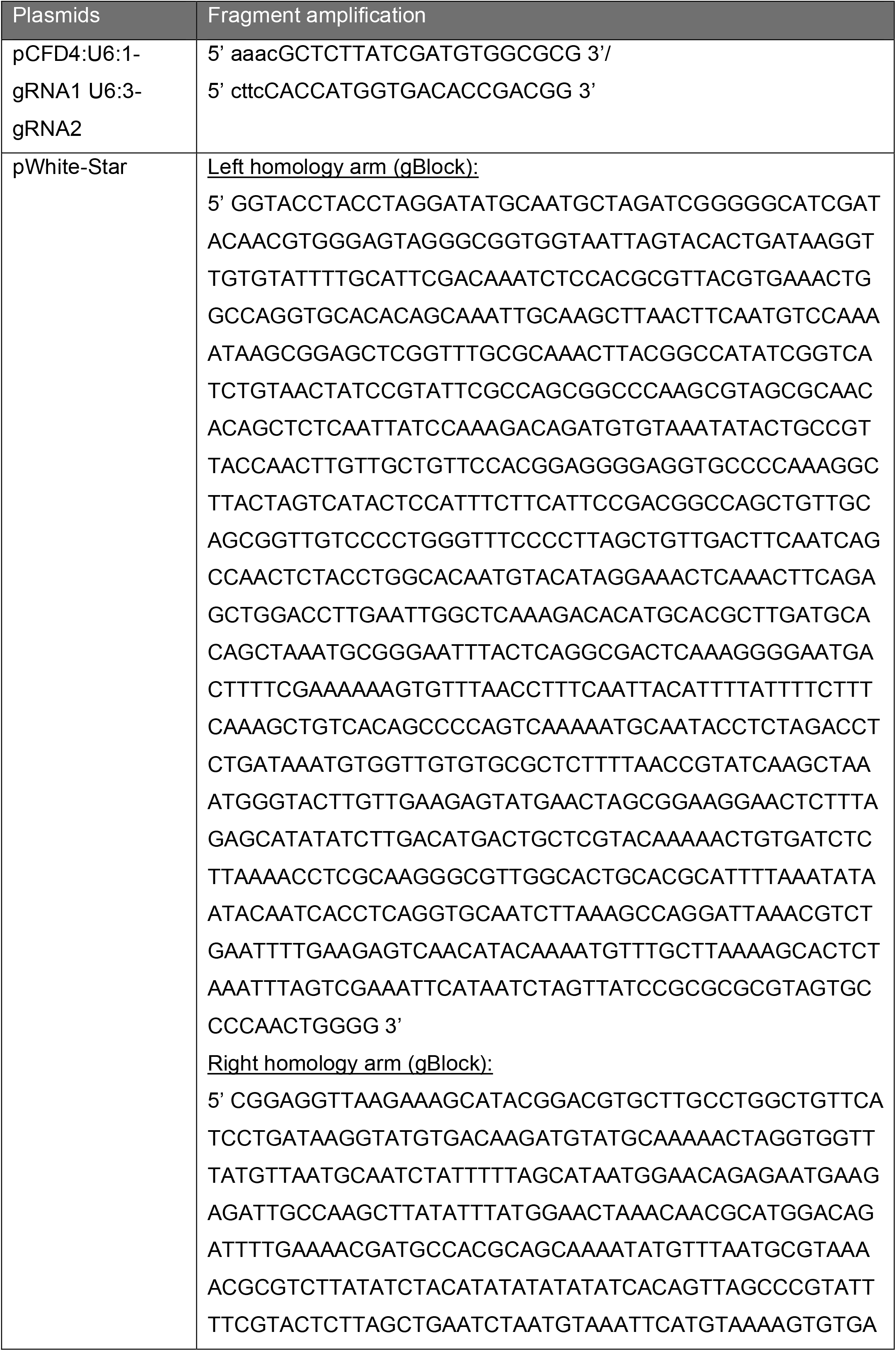
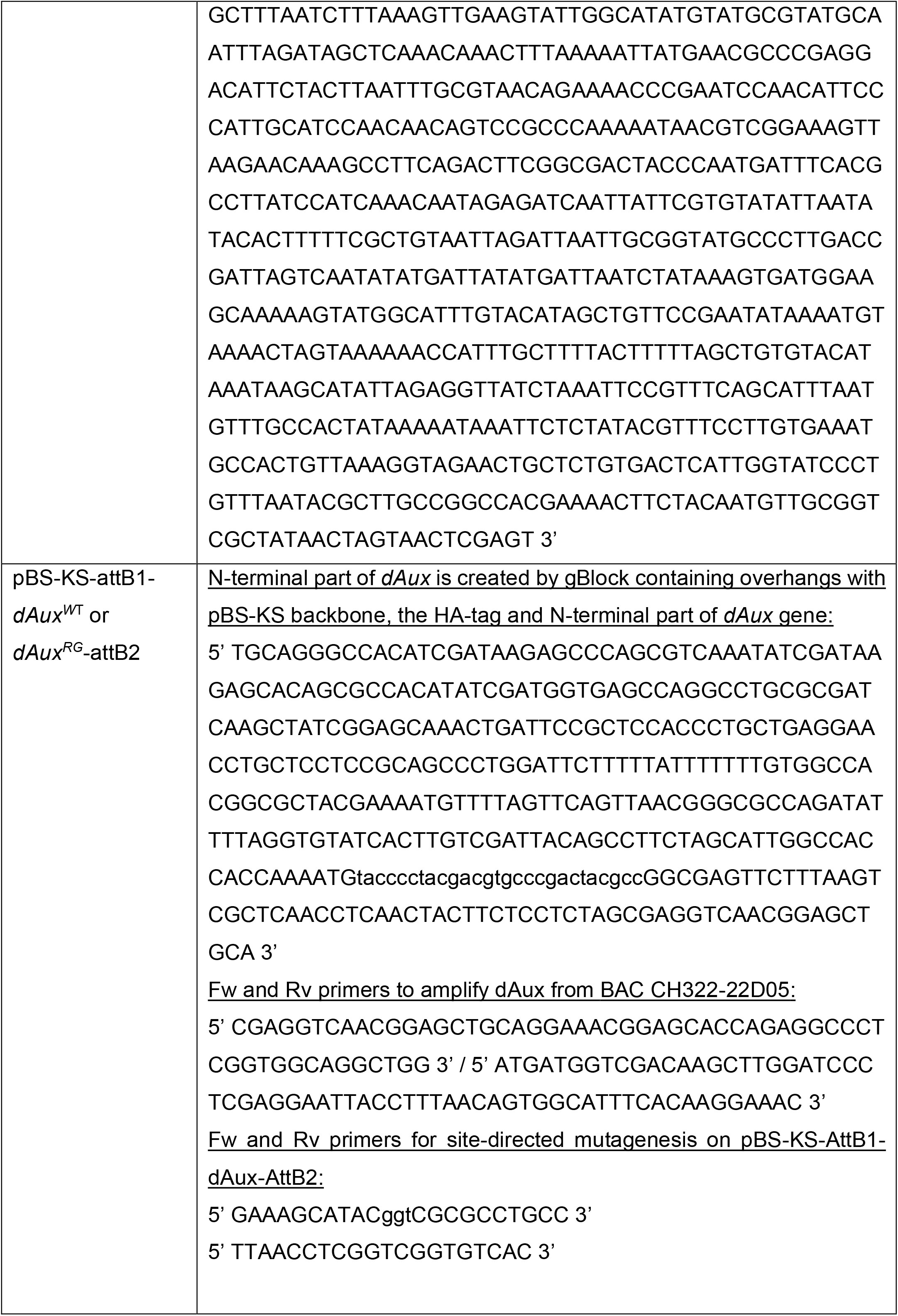
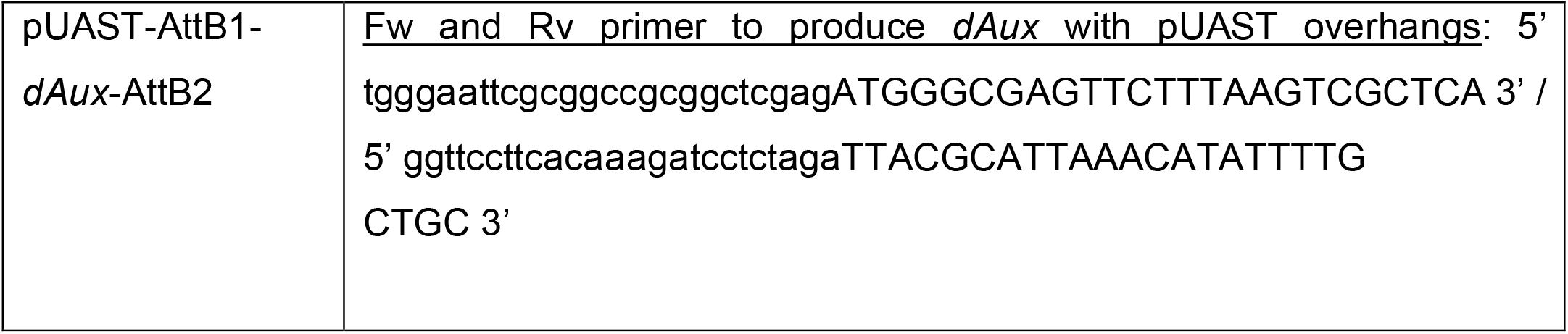

### Behavioral assays

#### Negative geotaxis

Groups of 5 males and 5 female flies were transferred into a empty fly vial with a line at the height of 4 cm from the bottom of the vial. Flies were tapped down quickly and were then given 30 s to climb past the 4 cm mark. Three trials were conducted for each set of flies and the average percentage of flies crossing the mark across three trials was calculated and presented (Benzer, 1973).

#### Seizure assay

Seizure assays were performed with groups of 5 males and 5 female flies that were transferred into transparent vials and stimulmated by vortexing of the vial for 10 s a maximum intensity. Numbers of active and flies that were unable to stand were quantified 5 s later. Graphs are presented in % of flies having seizures (Fischer *et al*, 2016).

### Fluorescent labeling

#### Immunohistochemistry on larval NMJs

Larval preparations were fixed for 20 min at room temperature (RT) with 4% formaldehyde (Sigma #47608) in 1x phosphate buffered saline (PBS; pH 7.4), next larval fillets were permeabilized with 0.4% PBX (TritonX-100 in 1X PBS). Tissue was blocked for 1h with 10% NGS in PBX and incubated overnight at 4°C with primary antibodies. After several washes, larval fillets were incubated with secondary antibodies for 2h in blocking solution at RT and washed with 0.4% PBX. Samples were mounted in Vectashield (Vector Laboratories). The following antibodies were used to label third instar *Drosophila* larvae: mouse α-Dlg [1:50 (DSHB; 4F3)], rabbit α-HRP [1:1000 (Jackson ImmunoResearch)], rabbit α-HA [1:200 (Cell Signaling Technologies; C29F4)], mouse α-Dynamin [1:50 (BD Biosciences, clone 41)]. Alexa Fluor 488-/Alexa Fluor 555-conjugated secondary antibodies (Invitrogen) were used 1:500.

#### Microscopy and image analysis

NMJs were imaged on a Nikon A1R confocal microscope through a Plan APO 60x A/1.20 Water Immersion DIC N2 lens or on a Zeiss Airyscan microscope at 63× magnification. NMJ’s were imaged at muscles 12/13 in segment A3. Image analysis was performed using Image J version 2.3.0 unless otherwise stated, by a researcher blind to genotype. At least three animals from independent crosses per genotype were examined in each analysis.

### Transmission Electron Microscopy

Third instar larvae were dissected in fresh Ca^2+^ free HL3 (110 mM NaCl, 5 mM KCl, 10 mM NaHCO_3_, 5 mM Hepes, 30 mM sucrose, 5 mM trehalose, and 10 mM MgCl_2_, pH 7.2; (Stewart *et al*, 1994)), nerves were cut and larvae subsequently incubated for 10 min in HL3 with 1.5 mM CaCl_2_ and 60 mM KCl. Multiple steps of washing with HL3 before fixation removed the non-internalized dye. Larvae were fixed in 4% paraformaldehyde (Laborimpex, 15714) and 2.5% glutaraldehyde (Polysciences, Inc, 111-30-8) in 0.1 M sodium-cacodylate buffer pH 7.4 (Merck, C0250-500g) at 4°C for at least 24h. The next day the larvae were osmicated in 2% osmium tetroxide (Laborimpex AGR1023) for 2h on ice. After staining in 2% aqueous uranyl acetate solution (EMS #22400) for at least 1.5h and dehydration in an ascending series of ethanol solutions, the samples were embedded in Agar 100 (Laborimpex, AGR1031) and cured at 60°C for 48h. Ultrathin sections (70 nm) were cut with a Dumont Diamond Knife on a Leica UCT ultra-microtome and collected on copper grids (Van Loenen, 01805-F) and imaged on a JEM 1400 transmission electron microscope (JEOL) at 80 kV with a bottom mounted camera (Quemasa; 11 megapixels; Olympus) running iTEM 5.2 software (Olympus). Statistical analyses were performed on larvae from 3 independent crosses per genotype. More than 5 NMJs per larvae were imaged.

#### Machine learning to detect synaptic vesicles at NMJ

Electron micrographs were first denoised using ImageJ and Tikhonov algorithm using following parameters; lambda = 2, iterations = 5, and sigma = 1.058. Fifty-five random cropped denoised EM images of 512 by 512 pixels were manually annotated using QuPath (Bankhead *et al*, 2017) and exported as labelled images using a custom Script. Cropped images and associated labels have been used to train a new model using StarDist and exported to be used with the StarDist ImageJ plugin (Weigert *et al*, 2020). A lower probability threshold was used to detect all synaptic vesicles inside a given bouton. The synaptic vesicles detected are highlighted as ROIs from which total amount and diameter were quantified. Three random tiles of 500 by 500 nm per bouton were examined. More than four boutons per animal per genotype were imaged.

### Toluidine blue staining

Adult fly heads were decapitated and immediately fixed in 4% paraformaldehyde (Laborimpex, 15714) and 2% glutaraldehyde (Polysciences, Inc, 111-30-8) in 0.1 M Na-Cacodylate buffer pH 7.4 (Merck, C0250-500g) for 2h at RT. Samples were further fixed at 4°C overnight, and then washed with 0.1 M Na-Cacodylate, pH 7.4, and subsequently osmicated with 2% osmium tetroxide (Laborimpex AGR1023). After staining in 2% uranyl acetate solution (EMS #22400) for at least 1.5h and dehydration in an ascending series of ethanol solutions, samples were embedded in Agar 100 (Laborimpex, AGR1031) and cured at 60°C for 48h. Alternatively, the sampes were kept substantially longer in 2% osmium tetroxide and kept overnight in 0.5% Uranylacetate/25% methanol solution before continuing with the dehydratation and embedding steps. Semi thin sections (1.5 µm) of the fly heads were collected on microscopy slides. The sections were then dried and stained on a heating block with a 1% toluidine blue (Merck, 89640-5G) solution including 2% Borax for 90 s at 60°C. The stained sections were mounted with Eukit Quick-hardening mounting medium (Sigma Aldrich, 03989-500ML) and imaged with the Leica DM25000M and 20x and 63x objectives. The percentage of normal vs abnormal ommatidia (defined as altered number and/or position of rhabdomeres) and the number of vacuoles in the fly retina were manually quantified using ImageJ.

### FM 1-43 dye uptake assays

Labelling of FM1-43 and quantification of intensities was performed as previously described (Verstreken *et al*, 2003). Briefly, third instar larvae were dissected in fresh Ca^2+^ free HL3 (Stewart *et al*, 1994), nerves were cut and subsequently incubated for 10 min in HL3 with 4 µM FM 1-43 (Invitrogen), 1.5 mM CaCl_2_ and 90 mM KCl. Multiple steps of washing with HL3 before imaging removed the non-internalized dye. NMJs were imaged at muscles 12/13 in segment A3 on a Nikon A1R confocal microscope through a 60X, 1.0 NA water immersion lens and stored using NIS elements software package. Mean boutonic intensities were determined, after background substraction, using ImageJ.

### Electroretinograms

Electroretinograms (ERGs) were recorded as previously described (Slabbaert *et al*, 2016). Briefly, flies were immobilized on glass microscope slides by use of 5 s fix UV glue. For recordings, glass electrodes (borosilicate, 1.5 mm outer diameter) filled with 3 M NaCl were placed in the thorax as a reference and on the fly eye for recordings. Responses to repetitive light stimuli were recorded using Axosope 10.7 and analyzed using Clampfit 10.7 software (Molecular Devices) and Igor Pro 6.37.

### Western Blotting

Fly heads were obtained from 30 flies from 3 independent crosses through flash freezing with liquid nitrogen, followed by vortexing. Homogenization of samples was obtained using a motorized pestle and 100 µL T-PER (Thermo Fischer Scientific) containing protease inhibitors (Sigma). 30 µg of protein was subjected to standard SDS-PAGE and Western blotting with HFP-conjugated secondary antibodies. Signal was visualized by West Pico Plus chemiluminescent reagent (Thermo Fischer Scientific) and an iBright imaging system (Thermo Fisher Scientific). The following antibodies were used: rabbit anti-GAPDH [1:2000 (Abcam)], (Pierce rabbit anti-DNAJC6 [1:1000 (Pierce)] and rabbit anti-HRP [1:5000 (Jackson ImmunoResearch)].

### Lipidomic Mass Spectrometry

Mass spectrometry was performed on heads of 15DO flies homogenized in 100 µl D-PBS (Dulbecco’s phosphate-buffered saline without Mg^2+^ and Ca^2+^) by Lipotype. Fifteen fly heads were pooled from three independent crosses for each analysis, and mass spectrometry was performed on duplicate samples (n=2). All liquid handling steps were performed using Hamilton Robotics STARlet robotic platform featuring the Anti Droplet Control for improved organic solvents handling. Prior to lipid extraction using chloroform and methanol, samples were spiked with lipid class-specific internal standards. After drying and resuspending in an appropriate MS acquisition mixture, lipid extracts were infused directly in QExactive mass spectrometer (Thermo Fisher Scientific) with TriVersa NanoMate ion source (Advion Biosciences). Samples are analyzed in both positive and negative ion modes, with MS resolution R_m/z=200_=280000 and MSMS resolution R_m/z=200_=17500, in a single acquisition. Acquired data was processed using lipid identification software based on LipidXplorer (Herzog *et al*, 2012). Data post-processing and normalization were performed by Lipotype using a developed data management system. Data analysis was performed using GraphPad Prism software 9.3.1. Pmol values (obtained by MS) of individual lipid species were transformed into a fraction of the total PA, DAG, PI, PC, PE, and PS lipids in the sample.

### Statistical analyses

Statistical analyses were performed using GraphPad Prism 9.3.1 software. Data is presented as mean ± SEM, unless otherwise stated. Differences between two groups were assessed using a two-tailed *t*-Test, differences between two or more variable were assessed by one-way or two-way ANOVA, with Tukey’s multiple correction test, unless otherwise stated. The criteria for significance are: ns (not significant), * p < 0.05, ** p < 0.01, *** p < 0.001, **** p < 0.0001.

## ACKNOWLEDGEMENTS

We thank the Bloomington Drosophila Stock Center for providing us flies and BestGene Inc for *Drosophila* embryo Injections. We thank the VIB Bio-imaging core Leuven and the Electron microscopy expertise unit at the Center for Brain & Disease Research. We thank Lipotype for the excellent service and the members of the Verstreken lab for helpful discussions and comments. We thank the lab of Dr. Henry. C. Chang for providing us the Aux^F956x^ fly line. This work was supported by FWO Vlaanderen, an ERC consolidator grant, the Chan Zuckerberg Initiative, a Methusalem grant of the Flemish Government, Opening the Future (Leuven University fund), Mission Lucidity to P.V. V.V. was supported by a Marie-Curie Sklodowska and an EMBO long-term fellowship and A.K. is supported by an FWO Vlaanderen fellowship. P.V. is an alumnus of the FENS-Kavli Network of Excellence.

## AUTHOR CONTRIBUTION

JJ, SK, VV and PV developed the project concept and designed experiments. JJ, SK and VV collected data. JJ and PV wrote the text and SK edited text. JJ, SK and PV analyzed data. JJ assembled all figures. BP and JS developed concepts for specific data sets, developed methods and/or collected data. JS processed all EM and histology samples. JJ, SK and PV interpreted EM experiments. JJ, SK, DC and AK collected and/or analyzed data related to ERGs. YW and NS designed constructs to make the new KI fly line.

## FIGURE LEGENDS

**Figure EV1.**
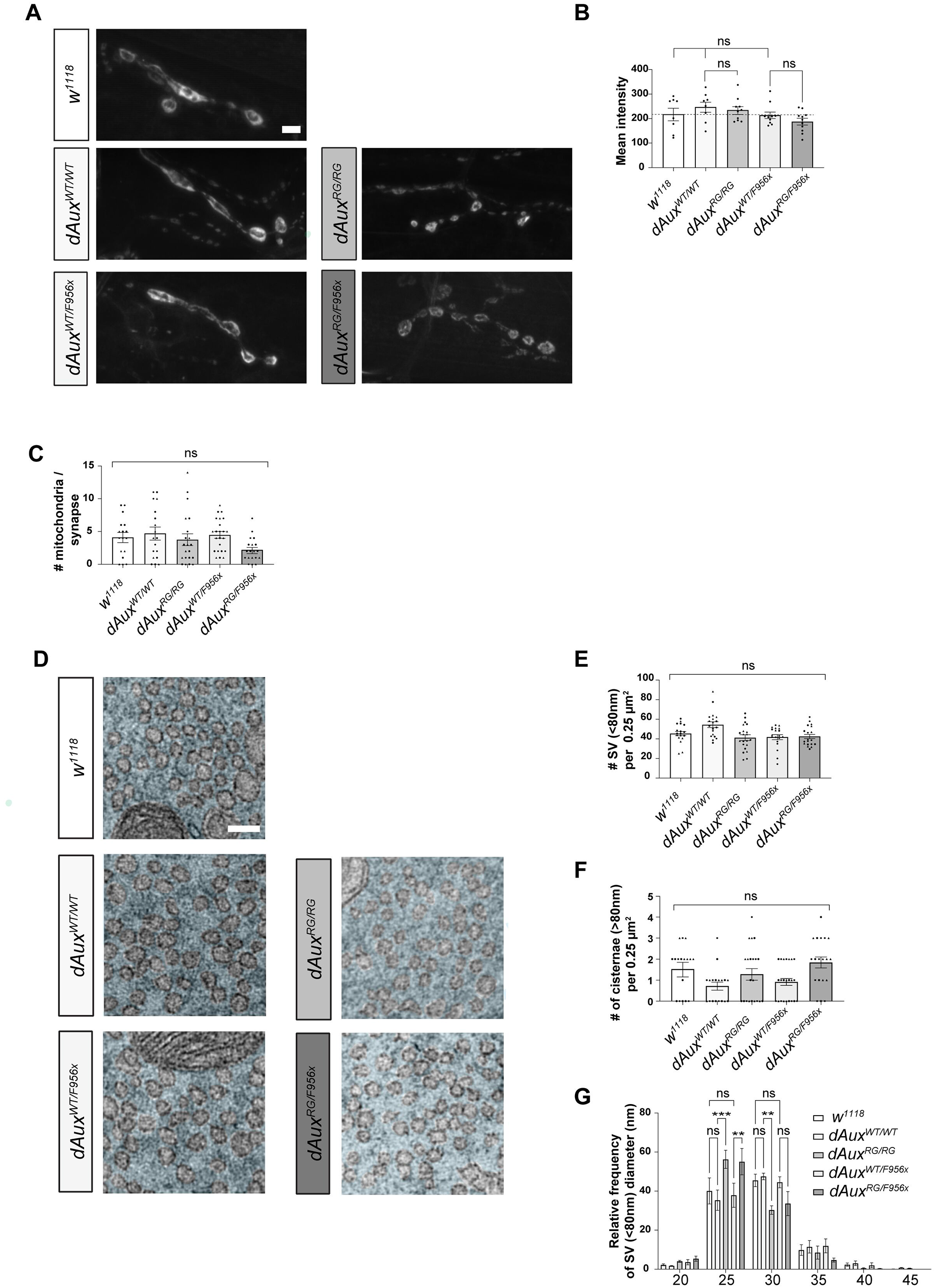
*dAux*^*RG/RG*^ or *dAux*^*RG/F956x*^ mutations do not obviously affect synaptic vesicle endocytosis at larval NMJs. A. Representative images of larval NMJs loaded (10min, HL3 with 1.5 mM CaCl_2_ and 90 mM KCl) in the presence of the lipophilic dye FM1-43 of the following genotypes: *w*^*1118*^, *dAux*^*WT/WT*^, *dAux*^*RG/RG*^, *dAux*^*WT/F956x*^ and *dAux*^*RG/F956x*^. B. Quantification of the mean FM1-43 intensity. Bars show mean ± SEM. Dotted line indicates mean of *w*^*1118*^. Dots represent mean intensity obtained from 5 NMJs per larvae. n≥8 L3 larvae (from independent crosses) per genotype were analysed. Bars show mean ± SEM. Kruskal-Wallis test with Dunnett’s multiple comparison test. ns: not significant. C. Quantification of the number of mitochondria inside each synapse. Bars show mean ± SEM. Dots represent the number of mitochondria per bouton and different shapes indicate the boutons belonging to the same animal. Kruskal-Wallis test with Dunnett’s multiple comparison test. ns: not significant. D. Synapse ultrastructure of NMJs of *w*^*1118*^, *dAux*^*WT/WT*^, *dAux*^*RG/RG*^, *dAux*^*WT/F956x*^ and *dAux*^*RG/F956x*^ larvae. Prior to TEM, larval fillets were incubated for 10 min in HL3 with 1.5 mM CaCl_2_ and 60 mM KCl. Scale bar indicates 100nm. E & F. Graphs represent the density of synaptic vesicles with a diameter below or above 80nm. Three random tiles of 500 by 500 nm (0.25µm^2^) per bouton were analysed by applying a machine learning algorithm (see methods). More than four boutons per animal and three animals per genotype were examined. Dots represent the average density calculated and different shapes indicate the boutons belonging to the same animal. Bars show the mean ± SEM. Kruskal-Wallis test with Dunnett’s multiple comparison test. ns: not significant. G. Synaptic vesicle diameter is shifted to lower values in boutons of *dAux*^*RG/RG*^ and *dAux*^*RG/F956x*^ mutants compared to their proper controls. Diameters of SV were measured as mentioned in D & E and the relative frequency distribution of the diameter in bins of five was calculated. Bars show the mean ± SEM. Two-way Anova with Tukey’s multiple comparison test. ns: not significant, ** p < 0.01 and *** p < 0.001.

**Figure EV2.**
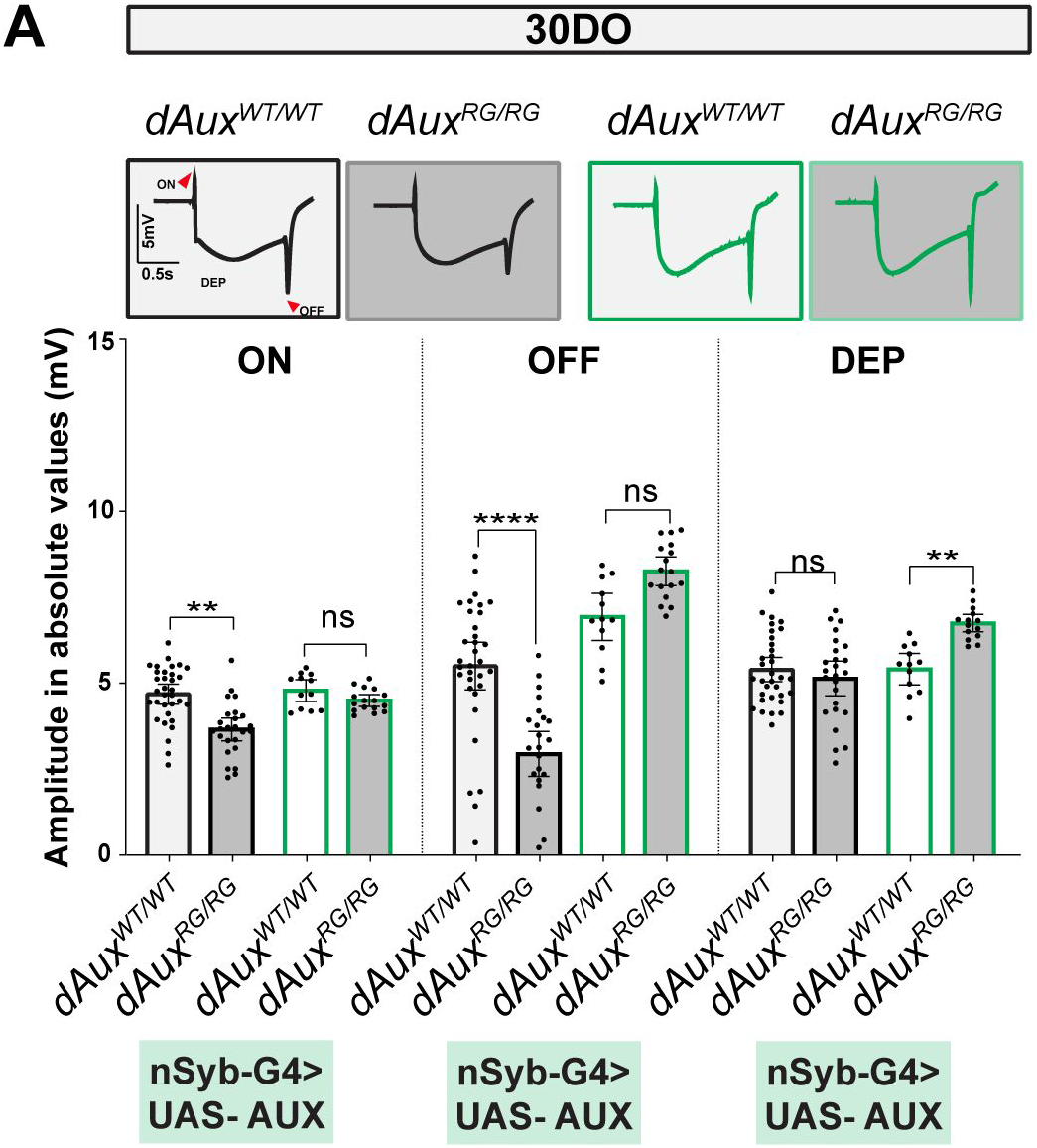
Neuronal expression of wild-type dAux rescues neuronal defects in 30DO *dAux*^*RG/RG*^ flies. A. Average ERG traces of 30DO *dAux*^*WT/WT*^ and *dAux*^*RG/RG*^ flies neuronally expressing Auxilin using the Nsyb-Gal4 driver. “ON” and “OFF” peaks are indicated with arrowheads. The ERG ON/OFF transient and amplitude of depolarization are quantified and represented as absolute values (mV) in the graphs below. Graphs represent the mean ± SEM of n≥10 per genotype, dots represent individual values. One-way ANOVA, Dunnett’s multiple comparison test. ns: not significant, ** p < 0.01 and **** p < 0.0001.

**Figure EV3.**
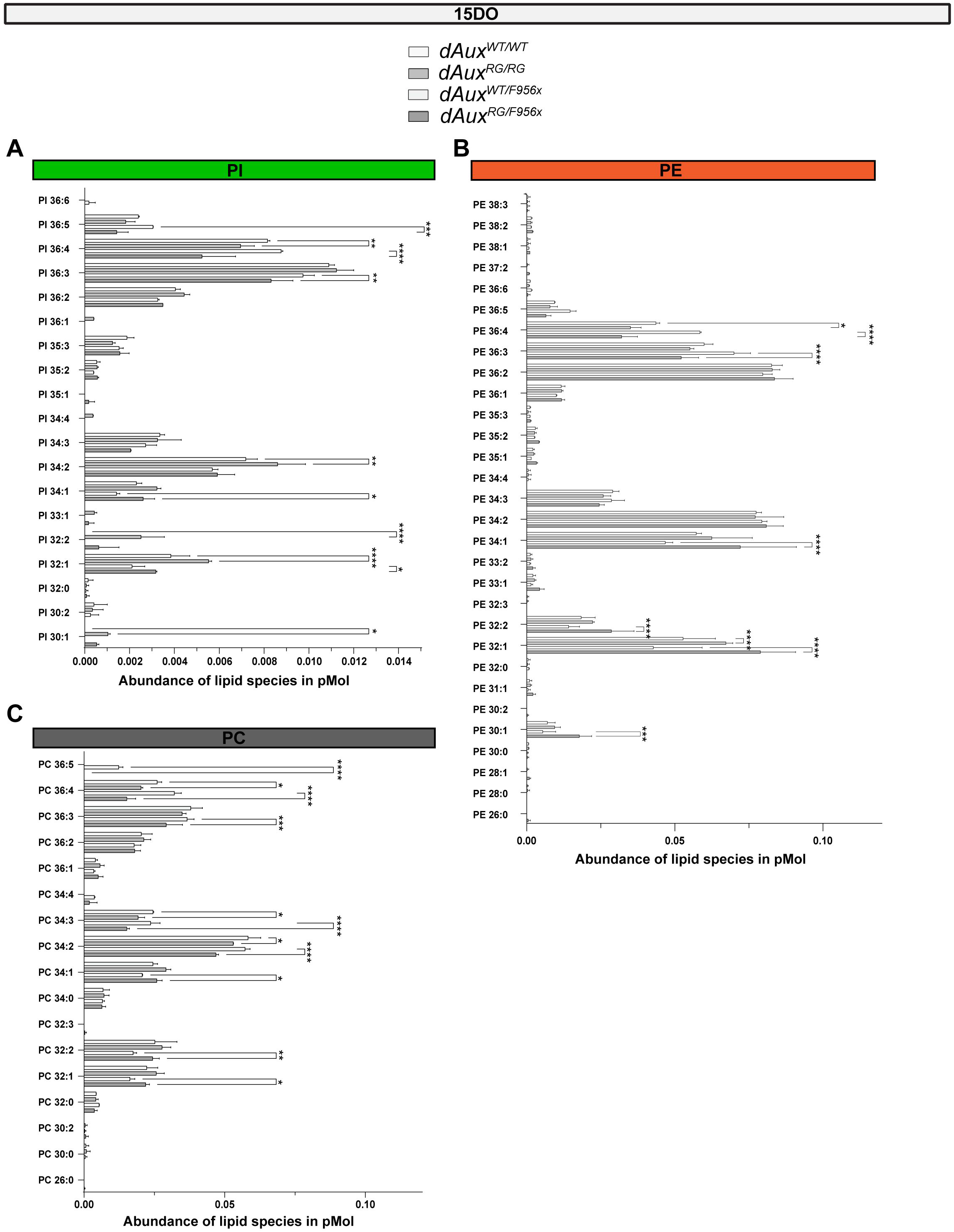
*Auxilin* mutant flies show alterations in PI, PE and PC species. A, B and C. The abundance of individual lipid species per GPL are presented in pMol. n = 2 MS analyses of 15DO fly heads collected from 3 independent crosses. Bars show mean ± SEM. Two-way ANOVA, Tukey’s multiple comparison test. *p<0.05, **p<0.01, ***p<0.001 and ****p<0.0001.

**Figure EV2.**
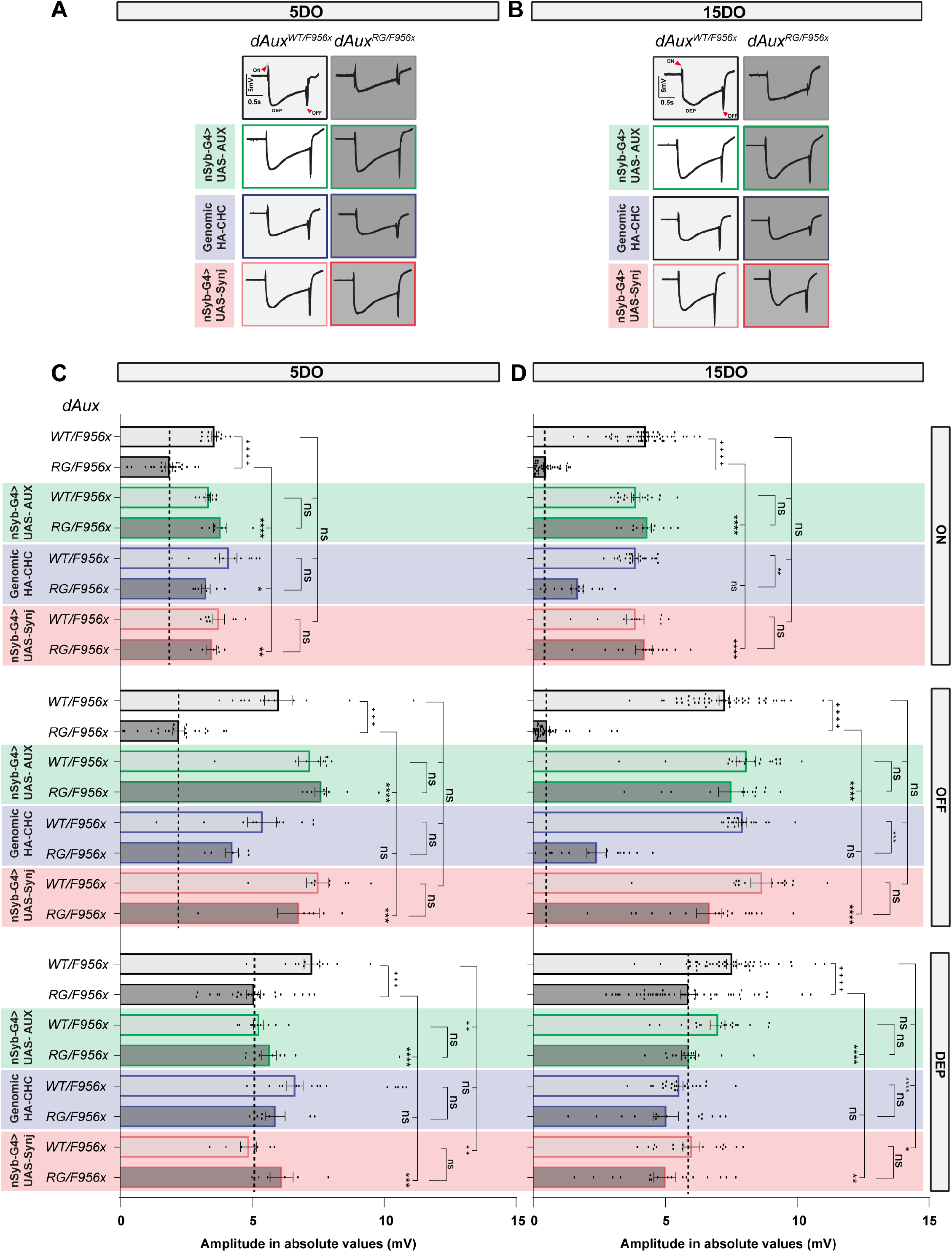
Expression of wild type Drosophila Synaptojanin restores neuronal function in Auxilin mutants. A & B. Average ERG traces of 5 and 15DO *dAux*^*WT/F956x*^ and *dAux*^*RG/F956x*^ flies that neuronally express wild type *Drosophila* Auxilin (UAS-AUX), harbor an extra genomic copy of wild type *Drosophila* clathrin heavy chain (HA-CHC) or neuronally express wild type *Drosophila* Synaptojanin (UAS-Synj). C & D. Quantification of the ERGs of 5DO (C) and 15DO (D) *dAux*^*WT/F956x*^ and *dAux*^*RG/F956x*^ flies that express wild type *Drosophila* Auxilin (UAS-AUX), harbor an extra genomic copy of wild type *Drosophila* clathrin heavy chain (HA-CHC) or neuronally express wild type *Drosophila* Synaptojanin (UAS-Synj).The ERG ON/OFF transient and amplitude of depolarization are quantified and represented as absolute values (mV). Graphs show mean ± SEM of n≥5 per genotype, dots represent individual values. One-way ANOVA, Dunnett’s multiple comparison test. ns: not significant, * p < 0.05, ** p < 0.01, *** p < 0.001, **** p < 0.0001.

